# The genetic basis of cone serotiny in *Pinus contorta* as a function of mixed-severity and stand-replacement fire regimes

**DOI:** 10.1101/023267

**Authors:** Mike Feduck, Philippe Henry, Richard Winder, David Dunn, René Alfaro, Lara vanAkker, Brad Hawkes

## Abstract

ABSTRACT

Wildfires and mountain pine beetle (MPB) attacks are important contributors to the development of stand structure in lodgepole pine, and major drivers of its evolution. The historical pattern of these events have been correlated with variation in cone serotiny (possessing cones that remain closed and retain seeds until opened by fire) across the Rocky Mountain region of Western North America. As climate change brings about a marked increase in the size, intensity, and severity of our wildfires, it is becoming increasingly important to study the genetic basis of serotiny as an adaptation to wildfire. Knowledge gleaned from these studies would have direct implications for forest management in the future, and for the future. In this study, we collected physical data and DNA samples from 122 trees of two different areas in the IDF-dk of British Columbia; multi-cohort stands (Cariboo-Chilcotin) with a history of mixed-severity fire and frequent MPB disturbances, and single-cohort stands (Logan Lake) with a history of stand replacing (crown) fire and infrequent MPB disturbances. We used QuantiNemo to construct simulated populations of lodgepole pine at five different growth rates, and compared the statistical outputs to physical data, then ran a random forest analysis to shed light on sources of variation in serotiny. We also sequenced 39 SNPs, of which 23 failed or were monomorphic. The 16 informative SNPs were used to calculate H_O_ and H_E_, which were included alongside genotypes for a second random forest analysis. Our best random forest model explained 33% of variation in serotiny, using simulation and physical variables. Our results highlight the need for more investigation into this matter, using more extensive approaches, and also consideration of alternative methods of heredity such as epigenetics.

## INTRODUCTION

Lodgepole pine (*Pinus contorta*) is very important conifer within the forests of Western Canada and throughout its range across much of Western North America. Ecologically, it is an important foundation species in the Rocky Mountain region, but it also constitutes a very valuable share of the region’s economy as a preferred source for both lumber and pulp (Coates & Sachs, 2012). Wildfires and mountain pine beetle (MPB) attacks have long been important contributors to the development of stand structure, particularly in the case of lodgepole pine. Recent studies have shown that MPB attacks and local fire regimes have a predictable effect on the structure of lodgepole pine stands (Alfaro *et al*., 2008; Alfaro *et al*., 2010; Axelson *et al*., 2009, 2010; Coates & Sachs, 2012; Hernández-Serrano *et al*., 2013; Scholefield, 2007). These studies have observed that stand-replacement (crown) fire regimes and infrequent MPB attacks tend to give rise to even-aged or single cohort pine stands, whereas mixed-severity (surface and crown fire) regimes and frequent MPB attacks produce multi-cohort (mixed age) stands. Jack pine has been shown to have a similar response to fire and MPB disturbance history (de Gouvenain & Delgadillo, 2012; Gauthier *et al*., 1996; Goubitz *et al*., 2004).

One of the adaptations that has made lodgepole pine so successful at regenerating after wildfire is the ability to produce serotinous cones. Serotinous cones are cones that do not open at maturity due to a resinous bond between the scales (Crossley, 1956; Lotan, 1967, 1976; Perry & Lotan, 1979; Radeloff *et al*., 2004), although serotiny is not reliably expressed in *P. contorta* until the tree has reached 50-60 years old (Perry & Lotan, 1979). When exposed to temperatures above 45°C, this resin melts, allowing the cones to open, and repopulate burnt-over stands with a sudden pulse of seed dispersal and recruitment (Cameron, 1953; Clements, 1910; Lotan, 1967, 1976; Perry & Lotan, 1979; Radeloff *et al*., 2004). *P. contorta* is not the only serotinous pine; there are at least 22 different serotinous species of *Pinus* including *P. canariensis, banksiana, P. halapensis, P. rigida* (Climent *et al*., 2004; Gauthier *et al*., 1996; Givnish, 1981; Habrouk *et al*., 1999; Lotan, 1976; Moya *et al*., 2012). Alternatively, some pines (including *P. nigra, P. monticola, P. albicaulis,* and *P. sylvestris*) do not have serotinous cones (Habrouk *et al*., 1999; McCune, 1988). Evidence has even been provided that seems to suggest that serotiny has evolved independently, multiple times within the genus *Pinus* (Grotkopp *et al*., 2004).

There have been multiple studies showing a relationship between cone serotiny and fire regime in pines. Givnish (1981) showed that pitch pine (*Pinus rigida*) stands with a history of high-frequency fire exhibited a higher occurrence of serotiny compared to those with lower frequencies of fire. Increased serotiny has also been demonstrated in areas of more intense (i.e. stand replacing) fires for *P. contorta* and *P. banksiana* (Gauthier *et al*., 1996; Radeloff *et al*., 2004; Schoennagel *et al*., 2003), and Schoennagel *et al*. (2003) showed that post-fire regeneration density was highest when pre-fire cone serotiny was highest. A number of other factors have since been shown to influence the proportion of serotinous cones in stands of lodgepole pine. Axelson *et al*. (2009) postulates that MPB attacks with only partial mortality of overstory lodgepole pine might favour non-serotinous cones, and thus lead to a multi-cohort stand. Benkman (2010; 2004; 2008; 2012) has shown that pine squirrels (*Tamiasciurus hudsonicus*) exert strong selective pressure against serotiny in lodgepole pine, to such an extent that the presence of pine squirrels can alter the stand structure over the course of several generations. Studies have also associated decreased serotiny with increased elevation (Tinker *et al*., 1994) and with increased latitude (Koch, 1996). In addition, a general trend of decreased serotiny has been observed with increasing longitude within the interior of British Columbia (Carlson, 2008).

Several studies have suggested that serotiny is under genetic control in *P. contorta* and *P. banksiana* (Lotan, 1967; Perry & Lotan, 1979; Rudolph *et al*., 1959; Teich, 1970) and that fire is likely a selective force for this trait (Climent *et al*., 2004; Gauthier *et al*., 1996; Givnish, 1981; Goubitz *et al*., 2004; Lotan, 1967; Parchman *et al*., 2012b; Perry & Lotan, 1979; Radeloff *et al*., 2004; Rudolph *et al*., 1959; Schoennagel *et al*., 2003). While early studies of serotiny generally considered it to be a binary phenotype (i.e. a tree is either serotinous or non-serotinous, and never partially serotinous) and under the control of just one or a very small number of genes (Lotan, 1967, 1976; Perry & Lotan, 1979; Rudolph *et al*., 1959; Teich, 1970), more recent evidence has shown that intermediate expression of serotiny does occur, and has suggested that serotiny is under the control of many genes, rather than just one or two (Parchman *et al*., 2011; Parchman *et al*., 2012a). Parchman *et al*. (2012b) analyzed 95,000 single nucleotide polymorphisms (SNPs), and identified 11 significant loci, the genotypes for which explained 50% of the phenotypic variation in serotiny, and the association remained consistent in each of 3 distinct populations of pines. This approach was the first published genome-wide association map of serotiny in pines and opened the door for a surge of new genetic analyses for serotiny in pines.

A better understanding of pine cone serotiny, its genetic basis, and its relationship to local fire regimes (and other environmental factors, such as the presence of seed predators) would further our understanding of the roles natural selection and genetic variation play in developing adaptive phenotypic variation (Parchman *et al*., 2012b). It could also have huge implications in the forestry sector. As climate change promises Western North America a sharp increase in the frequency, intensity, and severity of wildfires within the next decade (Budde *et al*., 2013; Flannigan *et al*., 2000; Fried *et al*., 2004; Running, 2006; Westerling *et al*., 2006), a better understanding of cone serotiny and the intricacies of all its interactions will prove invaluable in managing our pine forests, and ensuring they maintain the variability and adaptability that will allow them to cope. Beyond just (attempted) guiding of the response of lodgepole pine to these coming pressures, some sort of ability to predict the future response of lodgepole pine will cast broad implications on the future of our forests as a whole, as well as the entire forestry sector. This study could also serve as a resource for future investigations, which could potentially focus on assigning a more specific and quantitative description of fire regimes to better correlate potentially adaptive traits to fire as a selective agent.

We set out to examine and describe the sources of phenotypic variation of cone serotiny in *P. contorta*. To accomplish this, we constructed simulated populations of *P. contorta*, at 5 different growth rates (defined as an increase in population size). We then used a random forest approach to correlate various (simulated) population statistics and physical data from 122 trees of two separate fire regimes in central British Columbia with the observed serotiny of those trees. We also sequenced 39 SNPs from these trees, and ran another random forest analysis on the genotypes and calculated heterozygosity of the trees. Our approach yielded results that attribute a small amount of variation in serotiny to particular genotypes or other parameters, although confidence in these results is not high, partially due to the fact that 23 of the 39 SNPs failed or were monomorphic. Our results indicate that significant effort is still required in this area of research, and that alternative approaches to heredity (such as epigenetics) might yield better success in the future.

## METHODS

### SITE SELECTION

Data was collected from existing plots in six separate *P. contorta* stands that have previously documented and published disturbance histories and stand structures (Alfaro *et al*., 2008; Alfaro *et al*., 2010; Axelson *et al*., 2009, 2010).

Three stands from the Cariboo-Chilcotin area of British Columbia, near Alexis Creek and Tatla Lake (Figure 1) were examined. These stands are in the Interior Douglas-fir (IDF) biogeoclimatic zone, and the dry cool (dk) sub-zone. They have been previously shown by Axelson *et al*. (2010) and Daniels and Watson (2003) to be multi-cohort stands that have a mixed-severity fire regime and frequent mountain pine beetle disturbances.

**Figure 1.**
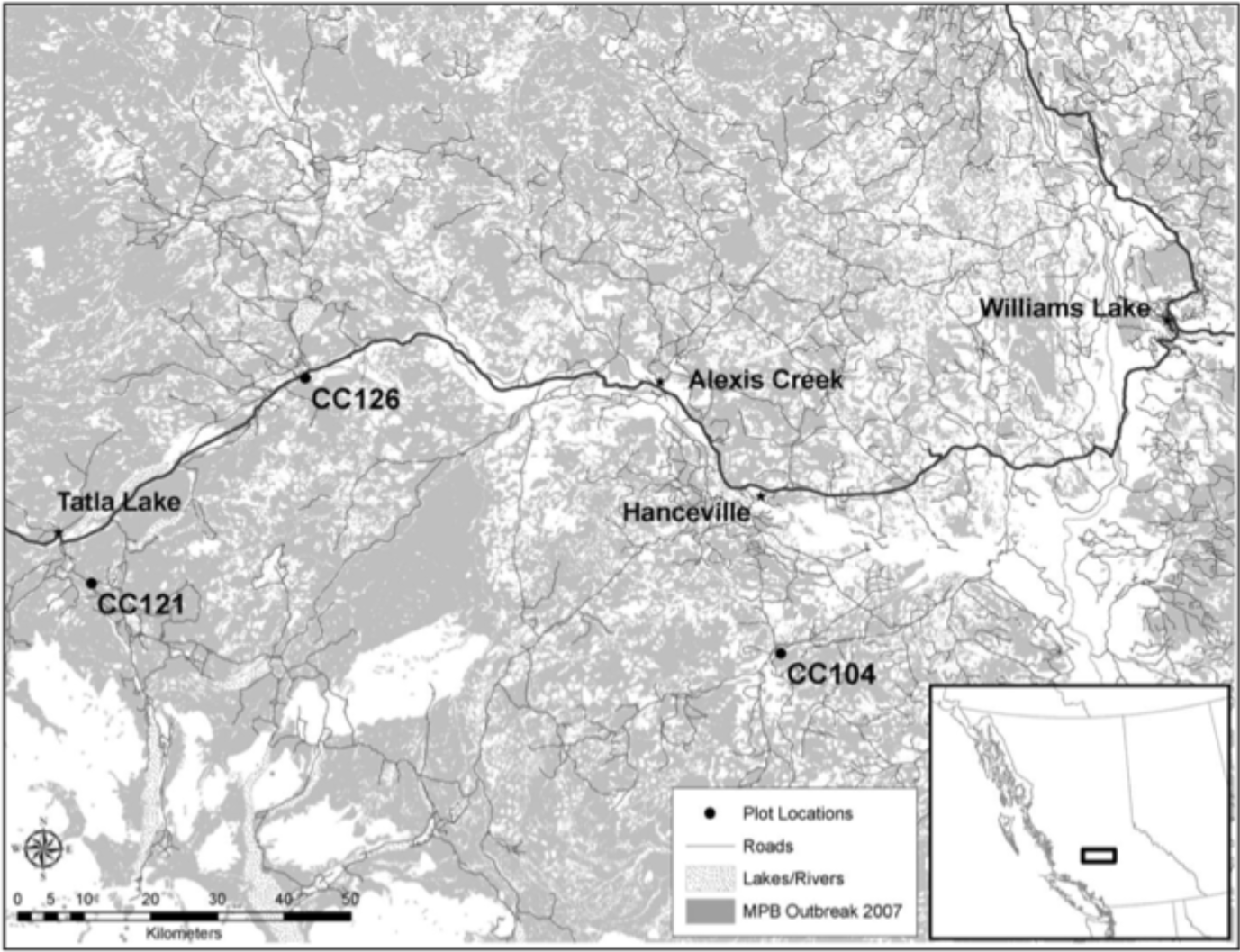
Geographic location of the three multi-cohort, mixed-severity fire regime stands of *P. contorta* from which needles and cones were sampled in B.C.’s Cariboo-Chilcotin. Adapted from Axelson *et al*. (2010).

The remaining three stands are in southern British Columbia, near Logan Lake (Figure 2). Axelson *et al*. (2009) described these as even-aged stands with a stand-replacing (crown) fire regime and infrequent mountain pine beetle disturbances. They also located in the IDF BEC zone, and the dk sub-zone. Unfortunately, one of the previously described stands has since been harvested, so it was replaced with an equivalent stand that has similar stand structure and disturbance history.

**Figure 2.**
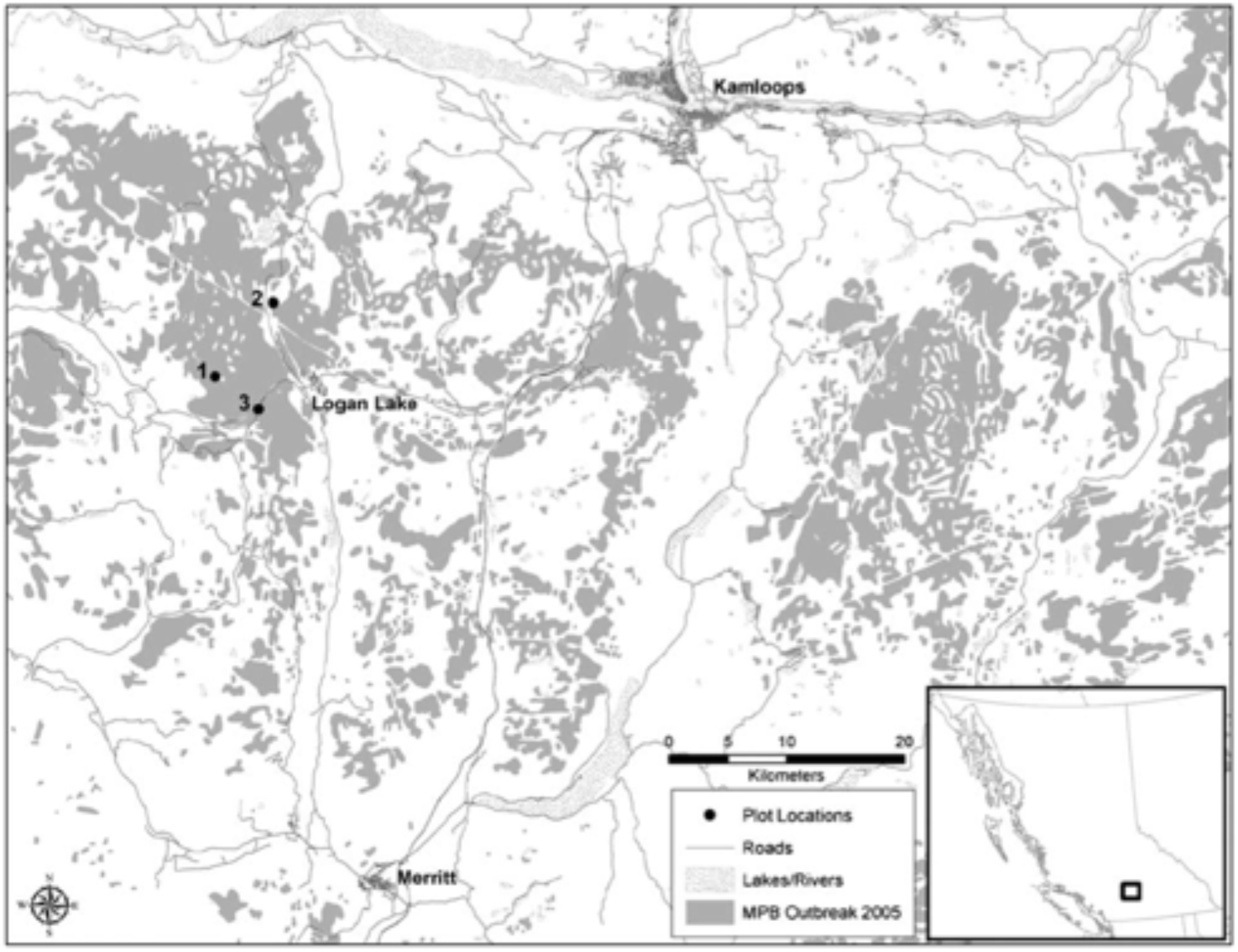
Geographic location of the three even-aged, stand-replacement fire regime stands of *P. contorta* from which needles and cones were sampled in B.C.’s southern interior. Adapted from Axelson *et al*. (2009).

### COLLECTION OF SEROTINY DATA

From each of the six stands, cone serotiny data was collected from twenty lodgepole pine trees. The trees chosen for sampling were healthy and greater than sixty years in age, to ensure full expression of serotiny. The live crown of each tree was visually divided into thirds (upper, middle, and lower), and two representative branches were selected from each third. With the aid of binoculars, the number of open and closed cones present on the distal 50 cm of each branch (six branches per tree) was recorded (in Logan Lake, 42 of the 60 trees were actually fallen, and cones were counted on the ground, rather than with binoculars). Trees with ≥75% closed cones were classified as serotinous. Additionally, 10-20 cones (a representative sampling) were collected from each of the 120 trees and left at room temperature in the lab for three weeks, during which they were monitored for opening. The percent of closed cones within these samples was recalculated, and averaged with that measured in the field to provide our ultimate measure of serotiny for each tree. Along with cone serotiny data, height-to-(live)-crown, diameter at breast height (DBH), and tree height were recorded for each tree.

### COLLECTION AND SEQUENCING OF GENETIC DATA

For each tree from which cone serotiny data was collected, needle samples were also taken. From each tree, we collected a minimum of 200 needles of the current year’s growth and stored them in a daily-refreshed desiccating environment. The needles from all 120 trees were prepared at the Pacific Forestry Centre before being sent to the McGill University and Génome Québec Innovation Centre, where their genotypes were sequenced for a set of 39 SNPs using SEQUENOM® IPLEX® Gold technology (Ehrich *et al*., 2005). Of these 39 SNPs, 11 were taken from Parchman *et al*. (2012a), and the remaining 28 from Cullingham *et al*. (2013b).

### ANALYSIS OF SEROTINY DATA

Serotiny data (serotiny, DBH, height-to-(live)-crown, and tree height) were compared in R (R Core Team, 2013) between Cariboo-Chilcotin and Logan Lake populations using side-by-side boxplots. A paired Wilcoxon test was then used to evaluate the difference of the means. A random forest analysis (a type of classification and regression tree analysis) (Breiman, 2001; Liaw, 2013) was run in R (R Core Team, 2013) using the physical field data alongside the statistical outputs of the simulated populations, looking for correlation to serotiny. This was performed for each of the 5 growth rates for the Cariboo-Chilcotin population and for the Logan Lake population.

### SIMULATION OF POPULATIONS

Populations were simulated using QuantiNemo (Neuenschwander *et al*., 2008) in Marlin (Meirmans, 2009). The simulation was run on 1000 individuals for 1240 generations, under logistic growth and random regulation models. The selfing (hermaphrodite) rate was set at 0.07 (Epperson & Allard, 1984; Perry & Lotan, 1979; Stoehr & Newton, 2002). A mutation rate of 1 × 10^−9^ mutations/site/year (Willyard *et al*., 2007) was used under a K-Allele model. The simulation was repeated for a range of growth rates (0.5, 1.0, 1.5, 2.5, and 5.0).

### ANALYSIS OF GENETIC DATA

The genotype data was used to calculate H_O_ and H_E_ for each sub-population using the adegenet package (Jombart, 2008; Jombart & Ahmed, 2011) in R (R Core Team, 2013). The values calculated for each sub-population were assigned to each tree in that sub-population.

Calculated heterozygosity values were calculated along with the genotypes of each tree/locus for explanation of serotiny using a random forest analysis (Breiman, 2001; Liaw, 2013).

## RESULTS

The top three explanatory variables (for serotiny) are shown for the random forest analysis of physical field measurements and simulation outputs (Table 1). DBH is one of the top three variables in each iteration. Tree height is present in the top three of each growth rate for Logan Lake, but in none for Cariboo-Chilcotin (Table 1). For the Cariboo-Chilcotin population, the percent of variation in serotiny explained by the best models is low or negative at growth rates of 0.5, 1.0, 1.5, and 2.5. At a growth rate of 5.0, however, the best model explains 33.3% of the variation in serotiny (Table 1). These values are higher in each of the Logan Lake growth rates, and all are positive. The highest value (19.23) is produced by a growth rate of 1.0 (Table 1).

**Table 1.**
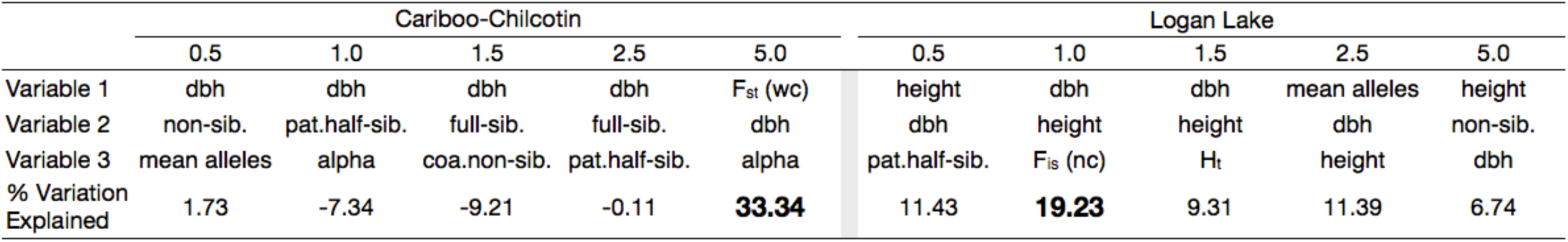
Top scoring explanatory variables for random forest analysis of physical field measurements and statistical outputs of simulated populations, repeated for each site, at each simulated growth rate. Also shown is the percent variation (serotiny) explained by the best random forest model of each analysis. Top variables include diameter at breast height [dbh], Weir & Cockerham (1984) Fst [Fst (wc)], tree height [height], mean # of alleles/locus/patch [mean alleles], mean proportion of non-siblings [non-sib.], mean proportion of paternal half-siblings [pat.half-sib.], mean proportion of full-siblings [full-sib.], between patch coancestry [alpha], coancestry of non-siblings [coa.non-sib.], Nei & Chesser (1983) FjS [FjS (nc)], and Nei & Chesser (1983) Ht [Ht]

Figure 3 shows the comparison of serotiny and physical field observations between the Cariboo-Chilcotin and Logan Lake populations. Logan Lake shows higher values than Cariboo-Chilcotin for height-to-crown, and overall serotiny, but lower for DBH, and similar for tree height.

**Figure 3.**
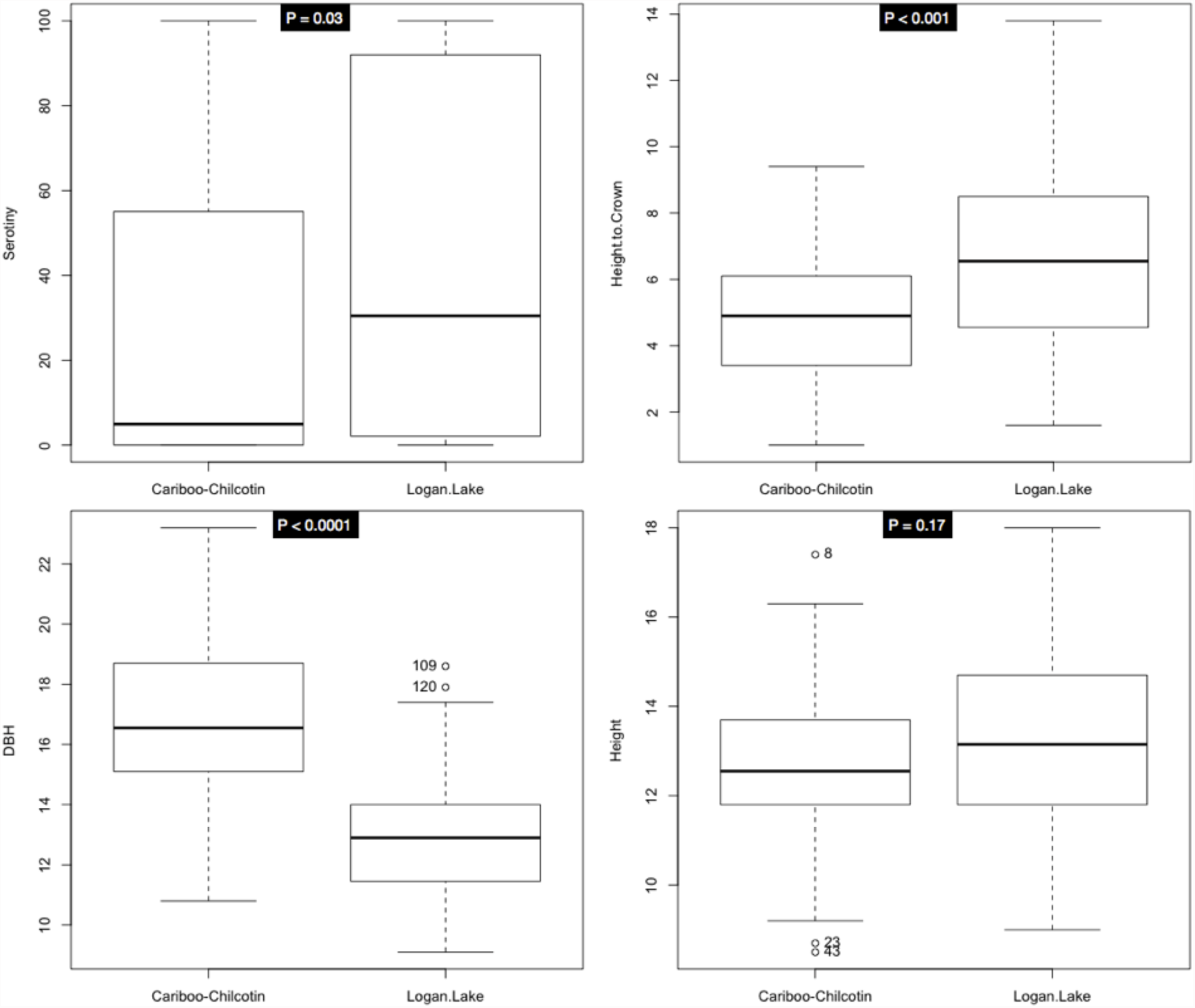
Side-by-side boxplots of serotiny, DBH, height-to-live-crown, and height of trees in Cariboo-Chilcotin and Logan Lake populations. Shown at the top of each plot is the p-value of a paired Wilcoxon test of the two means.

Tree height was the only comparison that showed a p-value (for a paired Wilcoxon test) of greater than 0.05 (p = 0.17) (Figure 3). The proportion of serotinous trees was higher in Logan Lake (28.3%) than in Cariboo-Chilcotin (16.1%) (data not shown).

Table 2 shows the results of SNP sequencing. Of the 11 SNPs developed by Parchman *et al*. (2012b), 6 failed, and the other 5 were monomorphic, leaving no descriptive SNPs (Table 2). Of the 28 Cullingham *et al*. (2013a), 10 failed and 2 were monomorphic, leaving 16 polymorphic SNPs on which further analyses were performed (Table 2). The sequencing results also showed that three trees from Cariboo-Chilcotin (one from stand 121, two from stand 126) failed to sequence and were not reported at all (not shown).

**Table 2.**
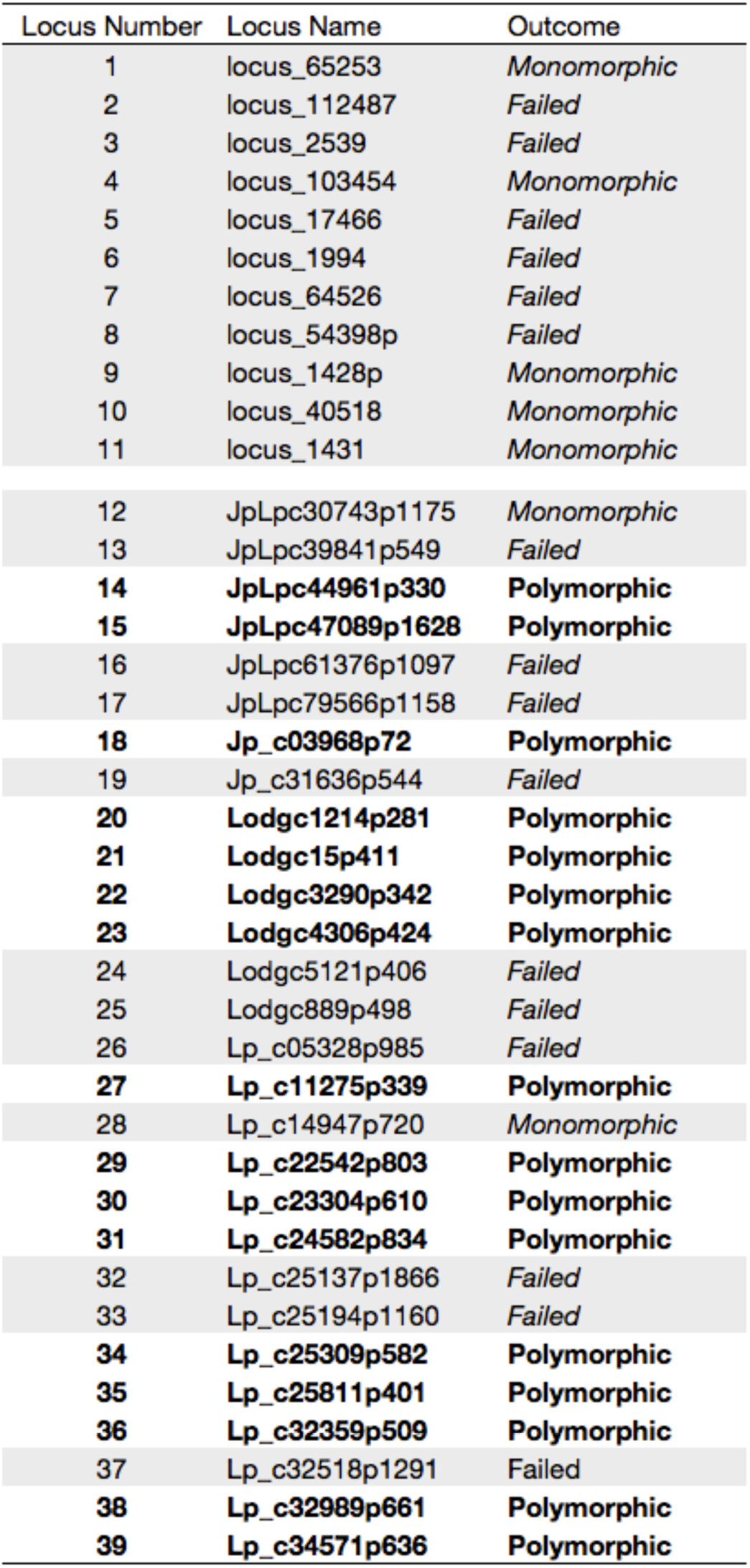
The outcome of Single Nucleotide Polymorphism (SNP) sequencing by McGill University and Genome Quebec Innovation Centre. Also shown is an arbitrary locus number for ease of reference. Loci 1-11 were adapted from Parchman *etal.* (2012) while loci 12-39 were adapted from Cullingham *etal.* (2013). Polymorphic SNPs were used for genotype analyses.

| Locus Number | Locus Name | Outcome |
| --- | --- | --- |
| 1 | locus_65253 | Monomorphic |
| 2 | locus_112487 | Failed |
| 3 | locus_2539 | Failed |
| 4 | locus_103454 | Monomorphic |
| 5 | locus_17466 | Failed |
| 6 | locus_1994 | Failed |
| 7 | locus_64526 | Failed |
| 8 | locus_54398p | Failed |
| 9 | locus_1428p | Monomorphic |
| 10 | locus_40518 | Monomorphic |
| 11 | locus_1431 | Monomorphic |
| 12 | JpLpc30743p1175 | Monomorphic |
| 13 | JpLpc39841p549 | Failed |
| 14 | **JpLpc44961p330** | **Polymorphic** |
| 15 | **JpLpc47089p1628** | **Polymorphic** |
| 16 | JpLpc61376p1097 | Failed |
| 17 | JpLpc79566p1158 | Failed |
| 18 | **Jp_c03968p72** | **Polymorphic** |
| 19 | Jp_c31636p544 | Failed |
| 20 | **Lodgc1214p281** | **Polymorphic** |
| 21 | **Lodgc15p411** | **Polymorphic** |
| 22 | **Lodgc3290p342** | **Polymorphic** |
| 23 | **Lodgc4306p424** | **Polymorphic** |
| 24 | Lodgc5121p406 | Failed |
| 25 | Lodgc889p498 | Failed |
| 26 | Lp_c05328p985 | Failed |
| 27 | **Lp_c11275p339** | **Polymorphic** |
| 28 | Lp_c14947p720 | Monomorphic |
| 29 | **Lp_c22542p803** | **Polymorphic** |
| 30 | **Lp_c23304p610** | **Polymorphic** |
| 31 | **Lp_c24582p834** | **Polymorphic** |
| 32 | Lp_c25137p1866 | Failed |
| 33 | Lp_c25194p1160 | Failed |
| 34 | **Lp_c25309p582** | **Polymorphic** |
| 35 | **Lp_c25811p401** | **Polymorphic** |
| 36 | **Lp_c32359p509** | **Polymorphic** |
| 37 | Lp_c32518p1291 | Failed |
| 38 | **Lp_c32989p661** | **Polymorphic** |
| 39 | **Lp_c34571p636** | **Polymorphic** |

In a random forest analysis of just the genotypes, the best model explained 9.88% of the variation in serotiny (Table 3 A). Of this 9.88%, Locus 27[CC] and Locus 14[CC] alone account for 7.02%. When the genotypes are combined with observed and expected heterozygosity (Table 3 B), the top random forest model explains 18.04% of the variation in serotiny. H_e_ at locus 35 and H_o_ at locus 30 have taken the first and second (respectively) scoring spots, while genotype CC at locus 27 is now third (Table 3 B).

**Table 3.** Top scoring explanatory variables for random forest analysis of **(A)** the SNP genotypes only, and **(B)** SNP genotypes as well as heterozygosity (observed and expected) values for each plot. Also shown is the percent variation (serotiny) explained by the best random forest model of each analysis.

|  | **A** |  | **B** |  |
| --- | --- | --- | --- | --- |
|  | Locus | Value | Locus | Value |
| Variable 1 | 27 | [ CC ] | 35 | He |
| Variable 2 | 14 | [ CC ] | 30 | Ho |
| Variable 3 | 30 | [ GT ] | 27 | [ CC ] |
| % Variation Explained | 9.88 |  | 18.04 |  |

## DISCUSSION

Our findings explain a small amount of variation in serotiny by simulated population statistics, physical (field) data, genotypes, and calculated heterozygosity, though these correlations are not significant enough to draw strong conclusions. Our results do suggest that the inheritance of serotiny is governed by a complex, multi-faceted set of loci. This contradicts the simple, one-gene framework previously suggested in the literature (Lotan, 1967, 1976; Perry & Lotan, 1979; Rudolph *et al*., 1959; Teich, 1970; Wymore *et al*., 2011), but seems to agree with more recent studies (Parchman *et al*., 2011; Parchman *et al*., 2012b). Our best model consisted of the simulated population statistics at a growth rate of 5.0 coupled with the field data from Cariboo-Chilcotin trees, accounting for 33.34% of the variation in serotiny. The strong preference for this (population) growth rate’s model over those of the other growth rates could suggest that the Cariboo-Chilcotin population is experiencing a similar rate of population growth. The same could be said of the Logan Lake population, with a growth rate of 1.0 (although the preference isn’t as strong). Across all models, DBH is a consistently high-scoring indicator of variation in serotiny in both populations. Linear regression analyses showed DBH to increase with increasing serotiny in Cariboo-Chilcotin, and have no correlation in Logan Lake (data not shown). DBH has been shown previously in the literature to have no significant effect on serotiny (Tinker *et al*., 1994), and should not be acting as a proxy for tree age, since all trees sampled were at least 60 years old, and therefore should be fully and reliably expressing any serotiny (Perry & Lotan, 1979). It is not clear why this pattern is observed here.

We found 20 genotypes that explained 9.88% of the phenotypic variation in serotiny, the top 2 of which were responsible for 7.02%. When combined with calculated heterozygosity values, the top 10 variables (6 genotypes and 4 H values) explained 18.04%. Again, confidence in the genotype results is low because the 11 SNPs that were supposed to be associated with serotiny (Parchman *et al*., 2012b) either failed or were monomorphic, and the 16 SNPs that were informative (polymorphic) have not been previously linked to serotiny in the literature (Cullingham *et al*., 2013b).

A large source of uncertainty in our results stems from the failure or lack of variability (monomorphic) in the genotyping of our 39 SNPs. Three trees (all from Cariboo-Chilcotin) failed to sequence at any of the loci, likely due to DNA degradation. Of the 39 SNPs, 16 loci failed and 7 were monomorphic. Most disconcerting was the fact that all 11 of the SNPs developed by Parchman *et al*. (2012a) that were expected to be related to serotiny came back non-informative (6 failed, 5 monomorphic). DNA degradation could be responsible, although it would more likely result in the failure of the entire sample from a particular tree failing (as was seen with the three trees that failed to sequences), rather than individual loci across all 119 samples. It is possible that the primers for these SNPs, while fixed in the populations from which they were developed, are actually variable in the populations from which we sampled (not universally fixed). It is also possible, though less likely, that the SNPs aren’t from lodgepole pine, and instead from some source of systematic contamination within the experiments for which they were screened. The presence of failures from both SNP sources (Cullingham *et al*., 2013b; Parchman *et al*., 2012a) would seem to favour the explanation of geographic variability, and SNPs that have been developed from primers that are fixed only in local populations.

Another shortcoming of this study was the small number of SNPs analyzed (39, compared to 95,000 in Parchman *et al*. (2012b)). A much larger set of SNPs would provide a better chance of capturing informative SNPs that are important to serotiny. With the development of new sequencing technologies, and the relative low cost of sequencing SNPs (Cullingham *et al*., 2013b), larger, more comprehensive genome-wide association studies are becoming increasingly feasible (Cullingham *et al*., 2013b; Parchman *et al*., 2012b). Thirdly, an inability to accurately, precisely, and quantitatively account for other factors that influence serotiny within our statistical models greatly limits their level of precision and ability to produce reliable results. Such factors include: presence of seed predators (Benkman & Siepielski, 2004), elevation (Tinker *et al*., 1994), latitude (Koch, 1996), longitude within interior British Columbia (Carlson, 2008), frequency and severity of MPB disturbances (Alfaro *et al*., 2008; Axelson *et al*., 2009), and possibly a threshold response to temperature among different trees (Parchman *et al*., 2012b).

Serotiny is a key adaptive trait in lodgepole pine, with significant influences in forest communities and ecosystems. As climate change brings about a marked increase in the frequency, intensity, and severity of wildfires for the foreseeable future (Budde *et al*., 2013; Flannigan *et al*., 2000; Fried *et al*., 2004; Running, 2006; Westerling *et al*., 2006; Wymore *et al*., 2011), a better understanding of cone serotiny and the factors that influence it will be an important tool for managing not just our pine forests, but our forests in general. Specifically, a genetic understanding of serotiny will be vital for the assisted migration and maintenance of variability and adaptability in pine forests as global climate change forces their displacement (Hoffmann & Sgrò, 2011; Malcolm *et al*., 2002; McLachlan *et al*., 2005; Scholefield, 2007; Winder, 2014). This study could also serve as a resource for future investigations, which could potentially focus on assigning a more specific and quantitative description of fire regimes to better correlate potentially adaptive traits to fire as a selective agent.

Our findings explain a small amount of variation in serotiny through correlation with various physical, genetic, and population statistic variables. More realistically, however, our results highlight the need for future investigation into this area, making use of more extensive and comprehensive association approaches as well as consideration of alternative (non-mendelian) methods of heredity such as epigenetics or maternal inheritance (Correns, 1937; Echt & Nelson, 1997). As technology advances and more and more knowledge is elucidated, quantitative descriptors of environmental factors shown to affect serotiny can be incorporated into statistical approaches, resulting in the progressive refinement of models and a finer limit of resolution in association studies.

## ACKNOWLEDGEMENTS

This work was part of Mike Feduck’s honours thesis in Biochemistry and Molecular Biology and was co-supervised by Philippe Henry and Brad Hawkes at UNBC.

Many thanks to Jennifer Anema for her work in preparing the genetic data, and to Jodi Axelson, Kevin Pellow, and George Dalrymple of the field team for helping to coordinate the trip, collect the field data, and for making my time with them thoroughly enjoyable.

## APPENDIX

**Figure 4a.**
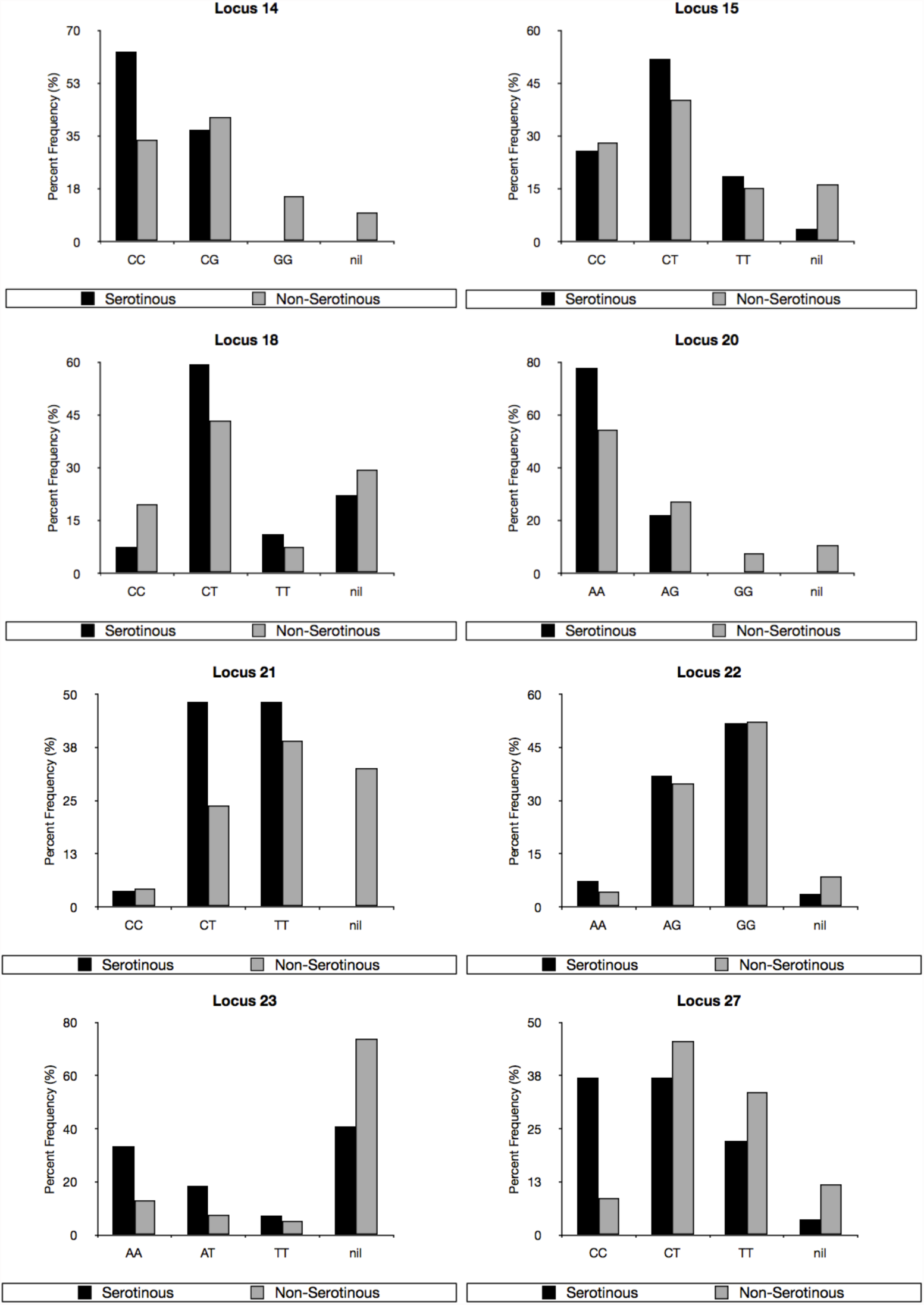
Observed proportions of each genotype among serotinous trees and non-serotinous trees. Loci 14, 15, 18, 20, 21, 22, 23, and 27 are shown

**Figure 4b.**
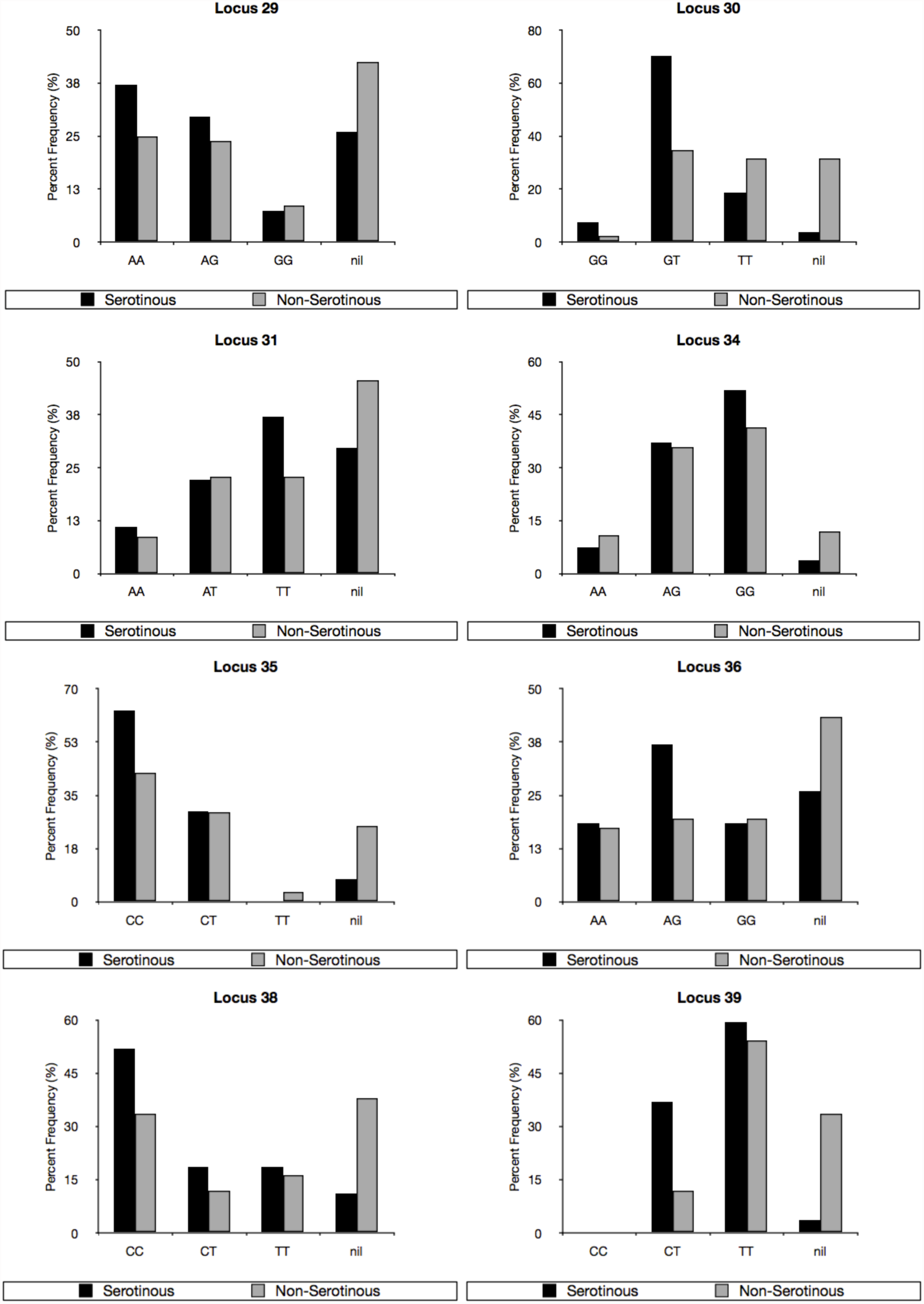
Observed proportions of each genotype among serotinous trees and non-serotinous trees. Loci 29, 30, 31, 34, 35, 36, 38, and 39 are shown

**Table 4.** Justifying the classification of serotinous and non-serotinous trees. Field and lab values for serotiny were averaged. Trees for which both field and lab observations indicated serotiny (≥ 75%) remained classified as serotinous (*). Four trees were classified as serotinous in the field, but after their cones opened in the lab, they were then classified as non-serotinous (**). These were classified as non-serotinous, attributing the ‘false’ classification of serotinous in the field to cones that were closed due to immaturity rather than serotiny. Two trees in the Cariboo-Chilcotin were classified as non-serotinous in the field, but serotinous in the lab (***). As open cones would not have closed overtime in the lab, the discrepancy was attributed to sampling errors in the cone collection, and the trees were classified as non-serotinous.

| Cariboo-Chilcotin |  |  |  | Logan Lake |  |  |  |
| --- | --- | --- | --- | --- | --- | --- | --- |
| Field | Lab | Average |  | Field | Lab | Average |  |
| 92.52 | 80.00 | 86.26 | * | 100.00 | 100.00 | 100.00 | * |
| 79.79 | 100.00 | 89.89 | * | 100.00 | 100.00 | 100.00 | * |
| 99.12 | 90.00 | 94.56 | * | 95.00 | 83.33 | 89.17 | * |
| 100.00 | 92.31 | 96.15 | * | 100.00 | 100.00 | 100.00 | * |
| 92.42 | 100.00 | 96.21 | * | 100.00 | 100.00 | 100.00 | * |
| 100.00 | 93.33 | 96.67 | * | 100.00 | 84.62 | 92.31 | * |
| 97.16 | 100.00 | 98.58 | * | 100.00 | 100.00 | 100.00 | * |
| 99.31 | 100.00 | 99.65 | * | 100.00 | 100.00 | 100.00 | * |
| 100.00 | 100.00 | 100.00 | * | 100.00 | 83.33 | 91.67 | * |
| 100.00 | 100.00 | 100.00 | * | 100.00 | 92.31 | 96.15 | * |
| 79.25 | 41.67 | 60.46 | ** | 100.00 | 100.00 | 100.00 | * |
| 76.00 | 66.67 | 71.33 | ** | 100.00 | 100.00 | 100.00 | * |
| 30.68 | 90.00 | 60.34 | *** | 100.00 | 100.00 | 100.00 | * |
| 56.52 | 78.57 | 67.55 | *** | 100.00 | 100.00 | 100.00 | * |
|  |  |  |  | 100.00 | 100.00 | 100.00 | * |
|  |  |  |  | 96.00 | 100.00 | 98.00 | * |
|  |  |  |  | 100.00 | 92.31 | 96.15 | * |
|  |  |  |  | 100.00 | 0.00 | 50.00 | ** |
|  |  |  |  | 100.00 | 0.00 | 50.00 | ** |

